# In vitro evolution of uracil glycosylase towards DnaKJ and GroEL binding evolves different misfolded states

**DOI:** 10.1101/2021.11.30.470542

**Authors:** Oran Melanker, Pierre Goloubinoff, Gideon Schreiber

## Abstract

Natural evolution is driven by random mutations that improve fitness. In vitro evolution mimics this process, however, on a short time-scale and is driven by the given bait. Here, we used directed in vitro evolution of a random library of Uracil glycosylase (eUNG) displayed on yeast surface to select for binding to chaperones GroEL, DnaK+DnaJ+ATP (DnaKJ) or *E.coli* cell extract (CE). Using binding to the eUNG inhibitor Ugi as probe for native foldedess, the CE selected population was further divided to Ugi binders (+U) (native) or non-binders (-U). We found that GroEL, DnaKJ and CE-U select and enrich for mutations causing eUNG to misfold, with the three being enriched in mutations in buried and conserved positions, with a tendency to increase positive charge. Still, each selection, as well as CE+U and natural evolution of eUNG has its own trajectory. While GroEL and CE-U selected for mutants highly sensitive to protease cleavage, DnaKJ selected for partially structured misfolded species with a tendency to refold, making them less sensitive to proteases. CE+U selected for more neutral mutations than natural evolution. In a more general context, our results show that GroEL has a higher tendency to purge promiscuous misfolded protein mutants from the system, while DnaKJ binds mutants misfolding-prone species that are, upon chaperone release, more likely to natively refold. CE-U shares some of the properties of GroEL and DnaKJ selected populations, while harboring also unique properties, explained by the existence of additional chaperones in CE, such as Tig, HtpG and ClpB.

## Introduction

HSP70 (DnaK) and Hsp60 (GroEL) are ancient, highly conserved classes of molecular chaperones (Hartl et al., 2011) that act as polypeptide unfolding enzymes (Finka et al., 2016). Both can use the energy of ATP-hydrolysis to forcefully unfold stably misfolded structures in aggregated polypeptides and thereby assist misfolding-prone proteins to reach and transiently stay in the native state, even under stressful conditions inauspicious to the native state (Goloubinoff et al., 2018). As such, these chaperones can buffer less stable genotypic variations (Agozzino and Dill, 2018; Finka et al., 2016; Libich et al., 2015; Rebeaud et al., 2021; Rutherford, 2003; Tokuriki and Tawfik, 2009). In the case of Hsp70-Hsp40-Hsp110 (DnaK-DnaJ-GrpE in *E coli*), chaperone-assisted unfolding-refolding occurs via DnaJ-binding motives, enriched with bulky hydrophobic amino acids on the surface of misfolded structures in aggregated polypeptides (Rüdiger et al., 2001) and DnaK-binding motives, generally misfolded structures which, once unfolded, consist of five consecutive hydrophobic residues, flanked by positive charges in un-structured polypeptide segments (Rüdiger et al., 1997). In the case of Hsp60 (GroEL-GroES in *E coli*), chaperone-assisted unfolding-refolding occurs via cooperative binding of exposed hydrophobic surfaces on aggregation-prone misfolded polypeptides, to a ring of hydrophobic motives displayed by the apical domains of the GroEL heptameric oligomers (Horovitz et al., 2022; Priya et al., 2013b). With the help of such chaperones, proteins can tolerate more destabilizing mutations, alleviating the evolvability penalty from loss of free energy. Yet, increasing the mutation load of a protein also increases its tendency to develop non-specific promiscuous interactions with other proteins in the very crowded cellular milieu, which is counter-productive. Because of their sticky nature, solvent-exposed hydrophobic residues are particularly problematic, and are thus constrained by purifying selection to maintain low hydrophobicity at a protein surface and reduce the likelihood of generating non-specific promiscuous sticky patches (Hochberg et al., 2020; Stiffler et al., 2015).

DnaK-DnaJ and GroEL preferentially interact with the misfolded states of proteins, with a strong preference for hydrophobic patches that are abnormally exposed to water. Following DnaJ-dependent interaction with substrate misfolded proteins and HSP70’s consequent ATP-dependent unfolding, the affinity of Hsp70 for the unfolded intermediate state was found to be significantly tighter than the affinity for the native state (De Los Rios and Barducci, 2014; Sekhar et al., 2018). A key aspect of the unfolding-refolding mechanism is that ATP hydrolysis adjusts the structure, and by this the relative affinity of the chaperone for the substrate towards a weaker affinity of the product (De Los Rios and Barducci, 2014; Rosenzweig et al., 2017). This reversibility is important because binding of first DnaJ, then of DnaK, or of GroEL oligomers, can inhibit aggregation, which following forceful unfolding and release of the unfolded polypeptide can at the same time favor the productive refolding to the native state. Acknowledging this multistep ATP-dependent binding-unfolding-release-refolding or re-aggregation mechanism of HSP60 and HSP70 chaperone activity, we aimed to determine the sequence and structural consequences of in vitro evolving a protein towards chaperone binding. For the *in vitro* evolution, we used the *E. coli* protein Uracil glycosylase (eUNG). This is a 229 amino acid protein, excises uracil residues from the DNA as a result of mis-incorporation of dUMP residues (Pearl, 2000). Folded eUNG binds with nanomolar affinity to the UNG inhibitor Ugi. Therefore, Ugi-binding can serve as a proxy for the structural integrity (native foldedness) of eUNG (assuming the incorporated mutations did not directly compromise binding). In vitro evolution was done by generating a library of randomly mutated Uracil glycosylase (eUNG-RL) proteins displayed on yeast surface, with FACS selection for binding to purified GroEL_14_ oligomers, DnaK+DnaJ2+ATP or to a total soluble cell extract from *E.coli*. Binding of Ugi was used as probe for eUNGs native foldedness (assuming that only folded eUNG will bind Ugi, see Figure 1). Both DnaKJ and GroEL were found to select for misfolded eUNG that did not bind Ugi, while the eUNG that were selected for CE binding, contained both an Ugi binding (+U) and non-binding (-U) population. Overall, the modulation of a protein evolutionary landscape orchestrated by these chaperones resulted, with evolutionary cycles, in the increasing accumulation of promiscuous mutants with non-native structures, suggesting that the DnaKJ and GroEL chaperones can salvage the proteome from mildly deleterious destabilization effects by some useful new mutations, while purging mutants with a dramatic propensity to aggregate, which can harm cellular functions.

**Figure 1.**
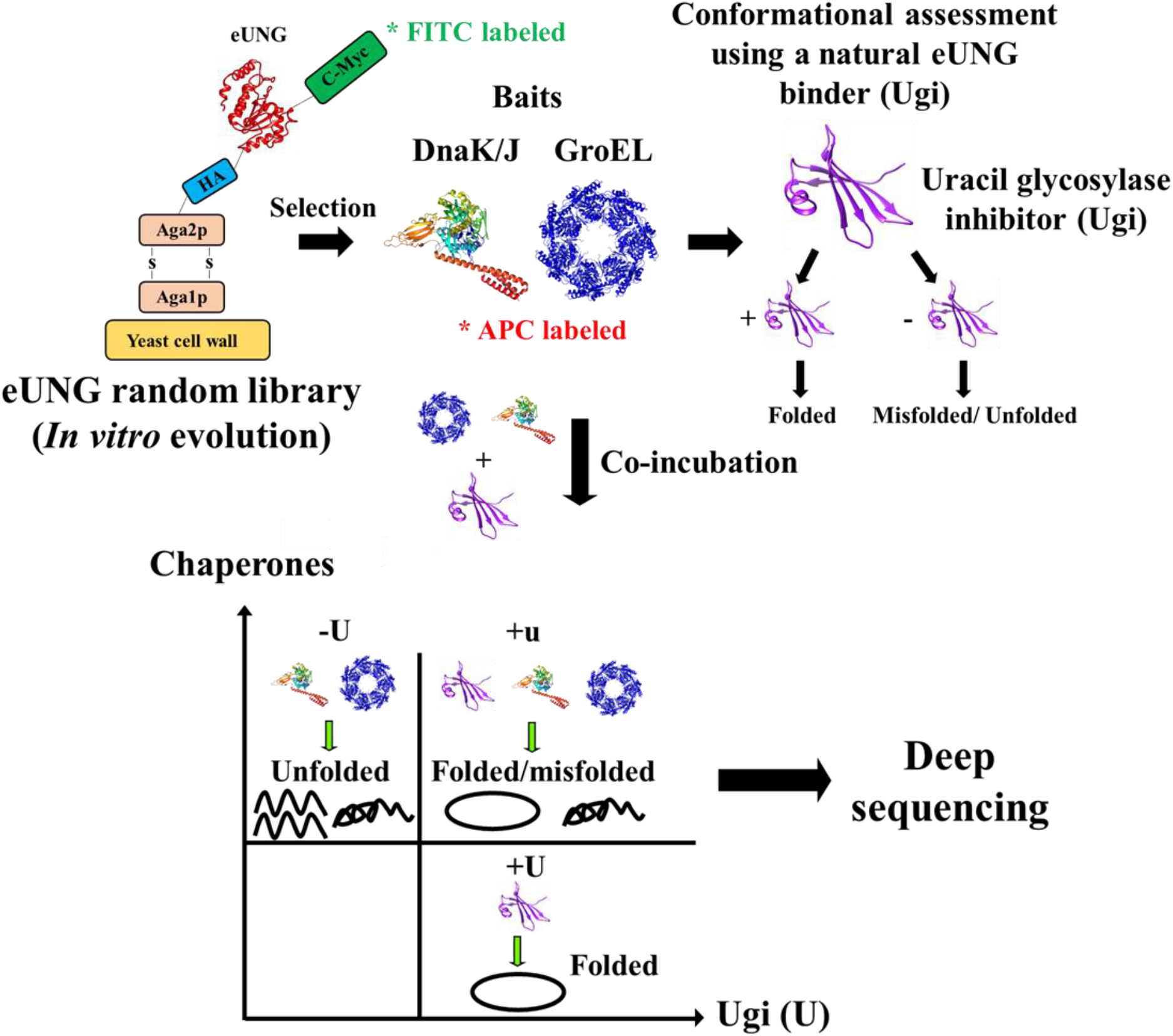
Experimental setup. A yeast surface display library of randomly mutated eUNG was prepared and selected against purified chaperones: DnaK+DnaJ+ATP (KJ) or GroEL-ATP. Folded and misfolded/folded variants were separated by their ability to bind fluorescently labeled uracil glycosylase inhibitor (Ugi). Selected clones binding Ugi are assumed to be folded, while those not binding Ugi are assumed to be misfolded or unfolded. Enriched populations were co-incubated with both chaperones and Ugi, to distinguish between folded and misfolded populations. We expect to see one populations, out of three possible, which binds only the chaperones, but not Ugi. eUNG variants from enriched populations were sequenced to study the relation between evolution of mutations and the baits used to enrich them.

## Results

### Evolving eUNG for chaperone binding

Chaperones have been shown to buffer evolution by increasing the allowed mutation load for marginally stable proteins, preventing their misfolding and aggregation (Rutherford and Lindquist, 1998; Tokuriki and Tawfik, 2009). Here, we examined the outcome of *in vitro* evolution of randomly mutated eUNG using chaperones as baits. First, we validated that externally supplied chaperones can specifically recognize and bind a protein displayed on the yeast surface when it is misfolded and not in its native conformation. Heat denatured fire-fly luciferase is known to bind DnaKJ and GroEL (Buchberger et al., 1996). Luciferase was inserted into the pCTcon2 yeast display vector C-terminal to Aga2p, where it is exposed to the surface, and incubated with pure GroEL_14_ oligomers or DnaK+DnaJ+ATP. These were incubated on ice, or were subjected to mild heat treatment (42°C) for 5 minutes, that is known to cause luciferase to irreversibly misfold in several minutes (without affecting the viability of yeast). We found that, as expected, the heat-denatured luciferase on the yeast-surface bound DnaKJ, and to a lesser extent GroEL, whereas native luciferase on ice did not bind the chaperones (Figure 2A). This experiment confirms that one can use yeast surface display to select for binding of misfolded proteins by chaperones. Interestingly, also *E.coli* cell extract did bind only heat treated luciferase and not luciferase on ice (Figure 2B), which may indicate that chaperones in the cell extract also bind mis-folded proteins, similar to that observed for pure chaperones.

**Figure 2.**
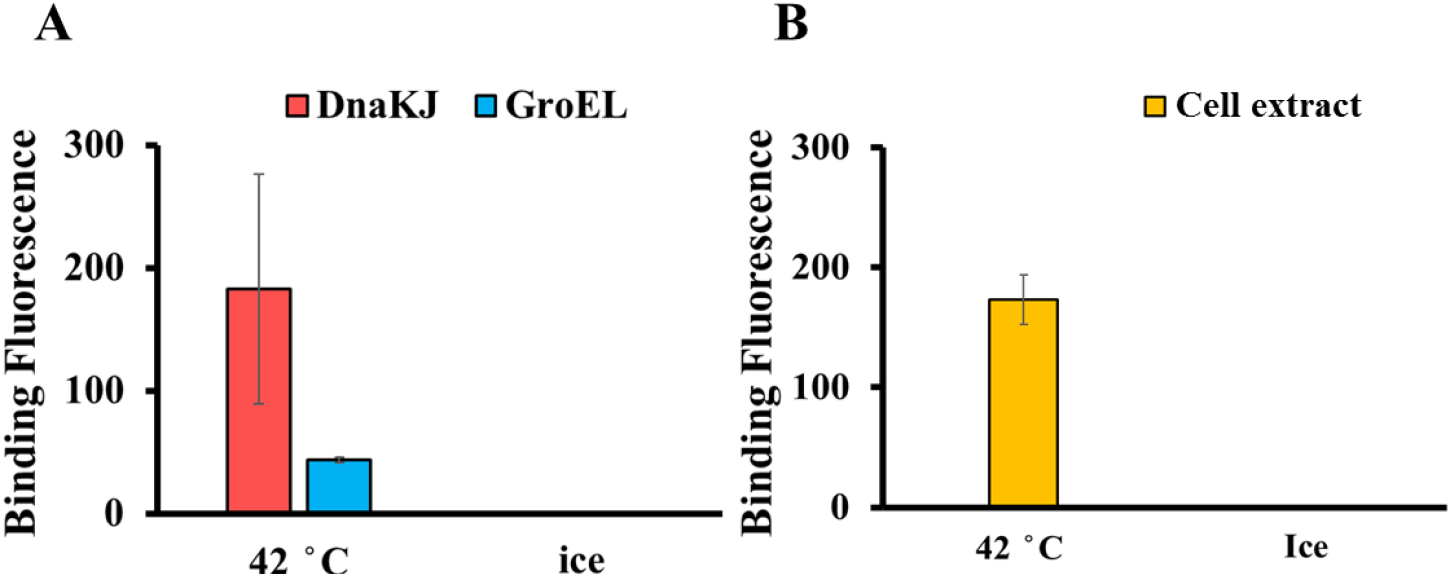
Chaperones bind mis-folded luciferase. **A.** Binding of labeled GroEL and DnaKJ to luciferase expressed on yeast surface. **B.** Binding of labeled cell extract (CE) to misfolded luciferase. For denaturation luciferase was incubated at 42 °c for 5 min with purified GroEL [1μM] and DnaKJ [2/0.4μM] proteins or cell extract [5mg/ml]. As controls, chaperones and cell extract were incubated with luciferase on ice, which keeps luciferase in a folded conformation. Binding fluorescence was analyzed by FACS. Controls were incubated on ice. Both chaperones were labeled with APC. GroEL conc. = 1uM, DnaK/J = 2/0.4uM. Y-axis - APC binding fluorescence (640nm) of normalized values.

### Evolving eUNG for chaperone binding

*In vitro* evolution of a randomly mutated protein is a powerful tool to study how the bait is dictating the fate of the evolved protein. Most commonly, the bait consists of another protein, where enhanced binding between the two is the outcome. Yet, being aware that mutation-induced misfolding of the bait can often produce non-specific promiscuous interactions with other proteins, here we choose to specifically address this by evolving eUNG-RL against purified DnaK+DnaJ+ATP (DnaKJ) or GroEL proteins and against a total cell extract. The random eUNG library was constructed by error-prone PCR, incorporating on average ~2-4 nucleotide substitutions per clone. The unsorted eUNG-RL underwent deep mutational scanning to determine the genetic complexity of the starting library. Using Enrich2 (see materials and methods) and Matlab (R2017a) the number of mutation-reads per position was extracted. Output reads were used to calculate the log-ratio of eUNG-RL (Table S1). The log-ratio is the common log of the total reads per position divided by the total number of reads including the number of WT reads normalized by the same calculation for the unsorted library using Eq. 1 (see materials and methods). Log ratio > 0 indicates a positive selection and log ratio < 0 a negative selection for a given position. Figure 3A shows the logo of eUNG-RL, and Figure 3B shows the enrichment of residue types of eUNG-RL relative to the theoretical calculated values, considering the propensity of amino acid changes upon single nucleotide mutations. This has been done as the frequencies of mutations requiring 2 nucleotide changes were very low, as expected. Probing the subset of amino acids reachable by single-nucleotide mutations is in line with natural evolution.

**Figure 3.**
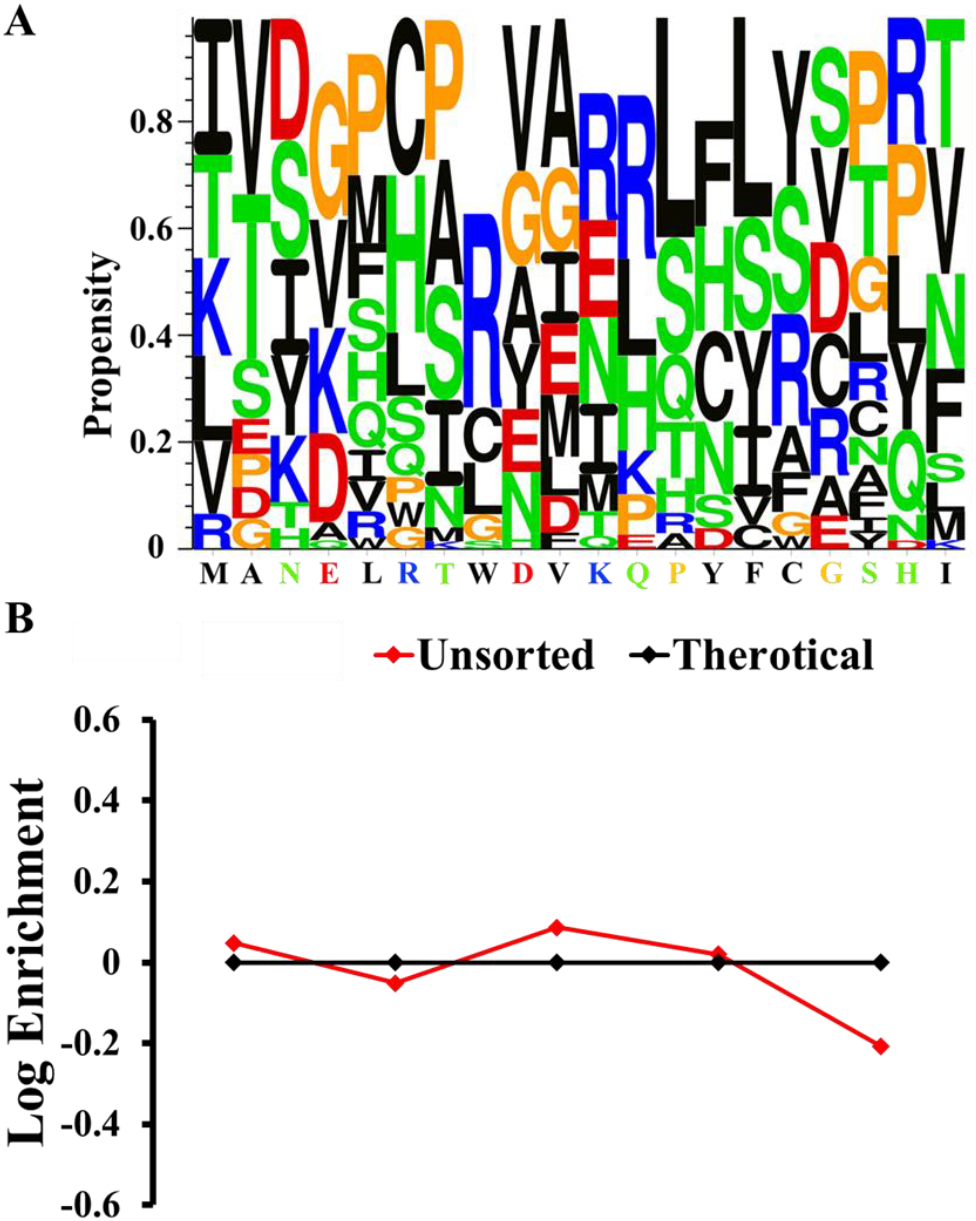
Natural evolution differs from *in-vitro* evolution in sequence space. **A.** Sequence logos display for the probability of WT amino acids to turn to any of 20 other amino acids in the unsorted library. Each WT amino acid and its corresponding possible mutation in the unsorted library was identified and its frequency was calculated as reads of possible mutations by the total number of reads for a possible mutations using Eq.4 where R_M_ denotes mutation reads over eUNG amino acid sequence. **B.** Hydrophobic, positive, negative, polar and other amino acids enrichment in the eUNG-RL (red) relative to the theoretical calculated values (black) assuming single-nucleotide mutations of the WT eUNG sequence. Analysis was done using MATLAB V.2017a

Having established that pure GroEL and DnaKJ (+ATP) bind preferentially to a yeast surface-exposed misfolded luciferase, but not to a natively-folded luciferase, and having established a random library of eUNG-RL, we next used DnaKJ and GroEL as bait against eUNG-RL. Each round of selection was evaluated by in vitro co-incubation of the yeast cells with the chaperones, to assess the type and degree of misfoldedness, and with Ugi to assess the native foldedness of the mutants. The progress of selection against DnaKJ and GroEL was found to be accompanied by a loss of binding to Ugi, indicating that the chaperones selected mostly for misfolded eUNGs (Figure 4A). Strikingly, the binding pattern for Ugi versus DnaKJ was mutually exclusive. Furthermore, tighter binding towards DnaKJ (with ATP) evolved more rapidly (after fewer evolution cycles) than binding towards GroEL (Without ATP), as judged by the increase in the APC signal. This suggests that the random library was selected for presumably misfolded eUNG–RL mutants with higher affinity towards DnaKJ, than towards GroEL. To confirm that the selection was not biased by the level of expression of different clones constituting the library, we compared the fluorescence values and distributions of cMyc expression between yeast displaying eUNG WT (without mutations) against our random library (Figure S1A) and found no difference between the two. Moreover, to avoid selecting for high-surface expressing clones (unfolded proteins may have lower surface expression), we gated for selection all c-Myc expressing clones in each round (Figure S1B). It should also be noted that the level of c-Myc expression was not constant with ongoing rounds of expression selected for misfolded proteins (Figure S1C), again verifying that we did not select for higher expression. In addition to selecting for GroEL and DnaKJ binding, we also selected for binding to *E.coli* cell extract (CE) (Figure S1D). Here, the selection resulted in two populations, one presumably native-like that did maintain Ugi binding and a second presumably misfolded that did not bind Ugi. These two populations were individually collected and called CE+U and CE-U.

**Figure 4.**
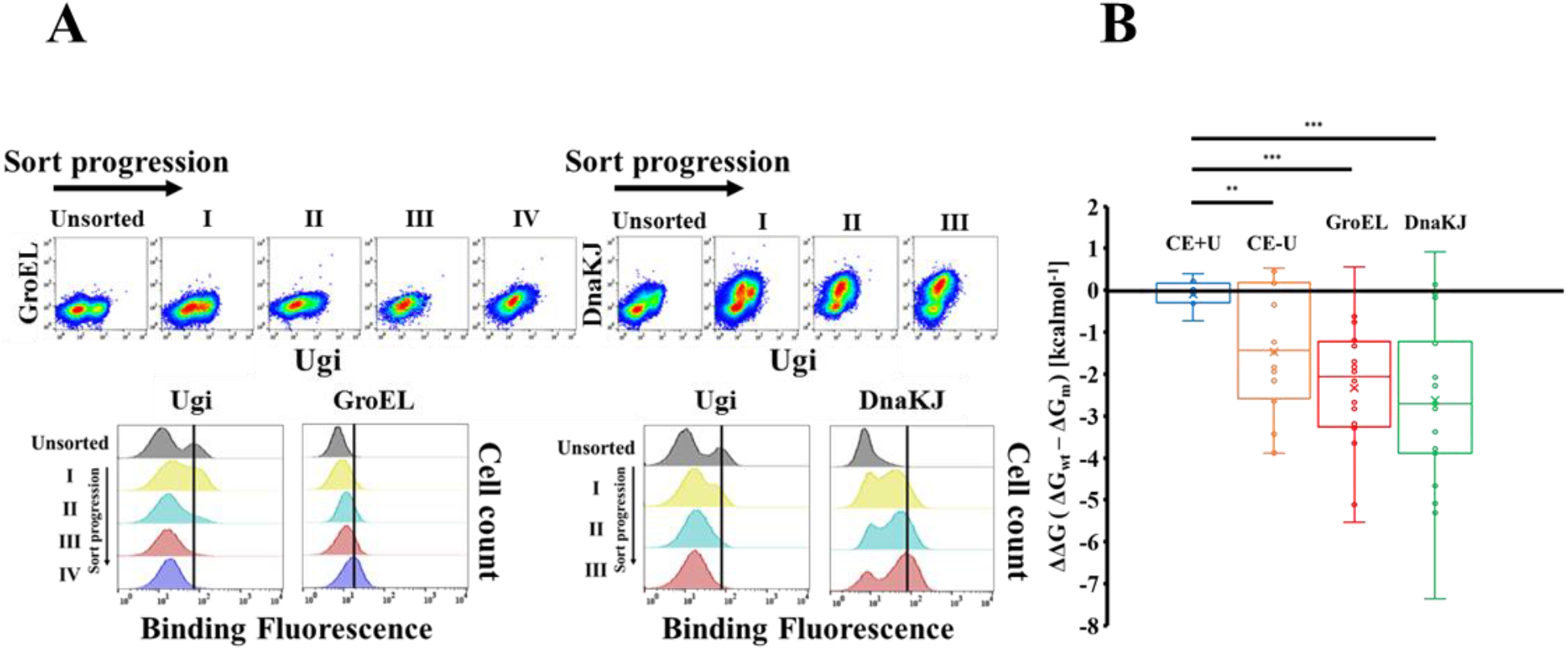
eUNG-RL selection against GroEL or DnaKJ results in loss of Ugi binding as a function of loss of free energy. **A.** eUNG-RL was selected against either GroEL (left) or DnaKJ (right), monitoring chaperon and Ugi binding after each round of selection. Progression of sort is indicated in roman numerals. Top, Scatter plots and bottom histograms of GroEL [1uM] and Ugi [5uM] as bait, binding yeast displayed eUNG-RL at each round of sort. After four rounds of selection, Ugi binding is not observed. Top right, scatter plots and histogram fit (bottom) of DnaKJ [2 or 0.4uM] and Ugi [5uM] with eUNG library at each round of sort. The gradual decrease in Ugi binding is visible. Black bar indicate maximum binding intensity for each substrate (chaperones or Ugi). Scatter plots: x axis - Ugi binding (FITC) and y axis - DnaKJ/GroEL binding (APC). Histograms: x axis - FITC/APC channel (530/640nM respectively) and y axis - cell count. DnaKJ was labeled by APC and Ugi was labeled by FITC. **B.** Free energy calculations for different sorted populations of eUNG. Box plot of the ΣΔΔG eUNG variants sorted against cell extract and chaperones as a function of +/- Ugi binding. CE+/-U stands for eUNG-RL selected against *E.coli* cell extract, with the clones selected as a function of binding Ugi and cell extract. ΔΔG was calculated using SDM and represents the difference in free energy between the eUNG wt and the mutant. Values represent the ΣΔΔG of mutation for a single variant. ** p-value < 0.01 *** p-value < 0.001.

### Thermodynamic characterization of selected eUNG clones as function of chaperone binding

To further investigate the degree of native foldedness of eUNG in the different selected populations, twenty single-clones from each population were isolated and sequenced. The mutational profile from each variant was used to calculate the change in free energy of folding of the individual mutations (ΔΔG [kcalmol^−1^]) (Worth et al., 2011), from which a ΣΔΔG was estimated for each variant (Figure 4B). For comparison, we also calculated ΔΔG values for CE+U and CE-U populations. Clones binding either GroEL, DnaKJ or CE-U had negative ΣΔΔG values (destabilized), with no significant difference between the GroEL- or DnaKJ-selected populations (Figure 4B). This is in strong contrast to the CE+U population, for which no change in calculated ΣΔΔG was found.

### Analyzing the mutational landscape of chaperone-sorted populations by deep sequencing

Mutations in the core of proteins or in conserved positions will more frequently translate in loss of folding free energy (Biol et al., 1992; Tokuriki et al., 2007). The degree in which a protein is destabilized changes the equilibrium between locally folded, transiently unfolded, stably unfolded (which is rare for globular proteins because of the tendency of exposed hydrophobic residues to avoid water), and stably misfolded conformations. Here, we examined the relation between the position and conservation of mutations in the different eUNG-sorted populations and their tendency to bind Ugi. Each sorted population and the unsorted eUNG-RL underwent deep mutational scanning and was evaluated as explained above, for the unsorted eUNG-RL. Output reads were used to calculate the log-ratio of each population (Table S1). Both GroEL and DnaKJ-selected population’s log-ratios showed reduced genetic complexity relative to the unsorted library (eUNG-RL). For mutations that were significantly higher than the mean of log-ratio > 0 (frequent mutations), the mean accessible surface area (ASA) and ConSurf conservation score (CS) were calculated (Table S2) (Ashkenazy et al., 2016; Krissinel and Henrick, 2007). The respective ASA and CS values of the different selected eUNG populations are shown in Figure 5A. As expected for a natively folded population, the mutations in eUNG sorted against CE+U were mostly surface-exposed and had a low conservation score. Conversely, mutations in eUNG sorted against either GroEL or DnaKJ were characterized by low ASA (buried) and high CS. The CE-U population resembles the chaperone selected population, suggesting it to be also unfolded or misfolded. Buried and conserved mutations increase the susceptibility of a protein to become non-native, explaining the loss of Ugi-binding in these populations. The results shown in Figure 5A are in line with those shown in Figure 4, with both suggesting that sorting against chaperones results in mutations that result in the protein being non-natively folded.

**Figure 5.**
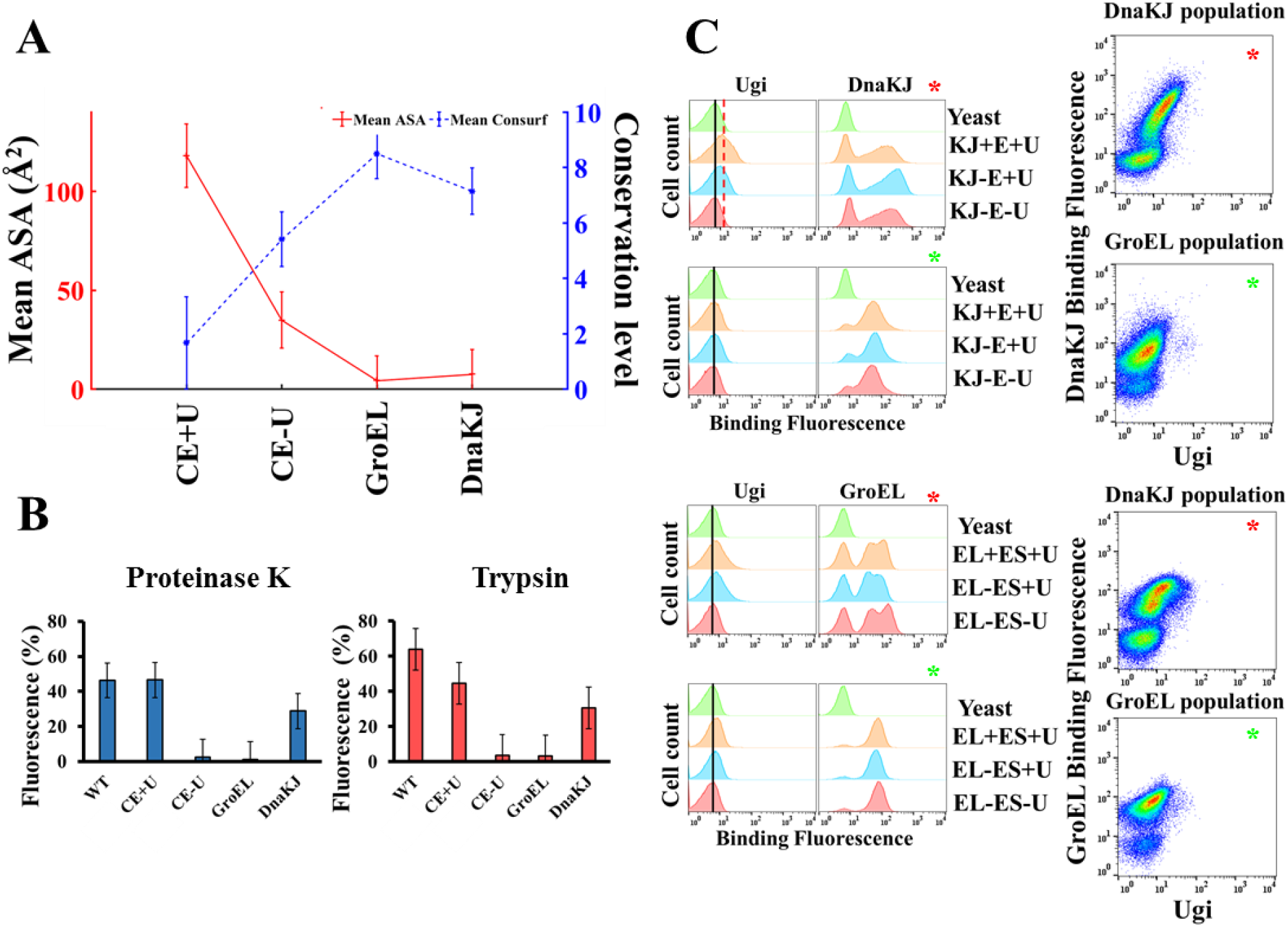
Biophysical properties of chaperone enriched populations. **A.** Negative correlation between accessible surface area (ASA) and ConSurf conservation score for CE, DnaKJ and GroEL selected populations. ASA and conservation scores were calculated using PDBePISA and ConSurf servers, respectively. Each selected population mean log ratio was calculated and positions with values significantly higher (p.value = 0.05 or 0.01) were used to calculate the mean ASA and ConSurf values. ASA is plotted with blue broken line and each population is marked with asterisk. ConSurf is plotted with red solid line and each population marked by a dot. **B.** Proteinase K (left) or Trypsin (right) treatment of enriched co-incubated populations. Each population was labeled with FITC and incubated with a limiting concentration of trypsin or Proteinase K, such that following 5 minutes at 37 ° C over 50% of the WT native eUNG resisted the protease treatment. Controls were incubated only with PBSX1. Change in fluorescence was analyzed using FlowJO and plotted as percentage of remaining fluorescence. **C.** Re-analysis of DnaKJ (panels 1, 3) or GroEL (panels 2, 4) selected population for DnaK [2μM]+DnaJ [0.4μM]+GrpE [1μM] +ATP (4mM) (panels 1, 2) or GroEL [1μM] GroES [1μM] +ATP [4mM] binding (panels 3, 4), respectively. Each was monitored also for Ugi [5μM] binding. Left panels show DnaKJ and GroEL selected population after refolding with their controls. KJ+/-E+/-U and EL+/-ES+/-U letters are initials for incubations with DnaKJ (KJ), GrpE (E), GroEL (EL), GroES (ES) and Ugi (U). Increase in Ugi binding affinity is depicted by the broken red line. Non-binding sub-populations are marked with black line. Right panels - scatter plots of DnaKJ and GroEL after refolding with DnaK+DnaJ+GrpE and GroEL+GroES.

### Susceptibility to protease cleavage of the sorted populations

Trypsin and Proteinase K are serine proteases that cleave after Arginine/Lysine or hydrophobic amino acids, respectively (EBELING et al., 1974; WALSH et al., 1964). Cleavage is dependent on the recognized site to be exposed, with buried sites being mostly shielded. Therefore, loss of native structure renders a protein more sensitive to digestion by a limiting concentration of a protease. Accordingly, chaperone-binding enriched populations should be more sensitive to protease digestion in accordance with their respective ASA, CS and their loss of Ugi binding. This hypothesis was examined by incubating enriched yeast populations with limiting concentrations of trypsin or proteinase K, for 5 min. The conditions were preset such that no more than 50% of the natively folded WT eUNG was cleaved. Protease digestion was measured by FACS as loss of FITC signal (due to yeast surface displayed eUNG being cleaved) and the percentage of remaining fluorescence was plotted. The CE+U binding population was almost as resistant to proteolysis as eUNG (Figure 5B). On the other hand, the GroEL and CE-U binding populations were completely degraded, in line with them being comprised of less compact unfolded and misfolded proteins. Surprisingly, different from the GroEL-selected population, the DnaKJ selected population was only partially digested, despite displaying the same reduction in ΣΔΔG (Figure 4) and high ASA and CS than the GroEL-selected population. This would suggest that the structural characteristics of the DnaKJ-selected population differs from that of GroEL-selected population.

To get a closer look at the difference between the GroEL and DnaKJ sorted populations, we took the last round of sort of both and re-analyzed their binding to DnaKJ or GroEL (Figure 5C). In addition, we also added GrpE and ATP+GroES, respectively to the DnaKJ and GroEL sorted eUNG. Whereas the co-chaperone GroES promotes the release from GroEL and the native refolding of the formerly GroEL-bound polypeptide substrate (Goloubinoff et al., 1989), GrpE functions as a nucleotide exchange factor promoting dissociation of ADP from the nucleotide-binding cleft of DnaK, and thereby driving substrate dissociation, which can lead to its native refolding. Figure 5C upper panel shows the DnaKJ-sorted population to have some residual Ugi binding characteristics, i.e. a potentiality to become native, particularly after the addition of GrpE. Conversely, the GroEL-sorted population had no Ugi binding activity, even in the presence of DnaKJ+GrpE or GroES+ATP (second and forth panels, respectively). The reciprocal experiment (third panel) shows that GroEL binds the DnaKJ-sorted population, however, in the absence of DnaKJ, no Ugi binding was observed. GroEL also bound, GroEL-sorted population, again without Ugi binding. Interestingly, GroEL bound two distinct DnaKJ-sorted populations (third panel), one stronger than the other, while the GroEL-selected population was homogeneous. This suggests that a more heterogeneous eUNG was selected by DnaKJ, in line with the limited proteolysis observed. This also suggests that sorting against GroEL results in mutations that are less compact and therefore with more exposed unfolded segments than sorting against DnaKJ. This is in line with our initial observation that heat-denatured luciferase binds DnaKJ better than GroEL (Figure 2A).

Previous reports have shown that both DnaKJE and GroELS recognize misfolded proteins and use energy from ATP hydrolysis to unfold and provide them with a new chance to reach the native state (Ben-Zvi et al., 2004; Priya et al., 2013a; Sharma et al., 2010), even under out-of-equilibrium conditions unfavorable the native state (Fauvet et al., 2021; Sharma et al., 2011). Yet, whereas GroEL can only unfold misfolded polypeptides which are smaller than 55 kDa and cannot disaggregate already formed proteins aggregates (Goloubinoff et al., 1989), DnaK can solubilize and unfold large polypeptides from already formed stable aggregates (Diamant et al., 2000). Our results further suggest that there is a fundamental difference between the two polypeptides unfoldases: while DnaKJE may target misfolding-prone protein mutants that can still be refolded to the native state, rendered metastable by the mutation, and thereby assist them to evolve into new native proteins, GroEL may target protein mutants that are more extensively damaged in their structure and contribute to their purging from the evolutionary process.

### Residue-dependent enrichment of selected populations

So far, we discussed the general properties of the different evolved populations (Figures 4 and 5). Next, we analyzed the mutation space of the evolved populations on a per-residue basis (Figure 6). The sequence space of the eUNG protein was divided by two parameters: 1. Log-ratio of the frequency to obtain a mutation at a given position of a sorted library divided by the frequency of the unsorted library (R0). 2. Frequency of a mutation in a specific amino-acid divided by the number of reads at the given position normalized by the unsorted library (Table S3). The first represents the spatial distribution (SD) of mutations per position in relation to the unselected library (positive or negative). The second, designates as specific spatial distribution – SSD, which describes the frequency of a specific position to be mutated into a specific amino acid (rather than any). Principle component analysis (PCA), which correlates population variance further verifies this conclusion (Figure 6A). PCA transforms a large set of variables into a smaller one, retaining most of the information content. PCA clearly identified the similarity between selections against GroEL, DnaKJ and CE-U in the main principle component axis (x-axis, 58%), while the three populations differ in the minor, y-axis. The unfolded populations are distanced from CE+U population. In addition, the log-ratio of SD of the different selected populations was clustered in hierarchal manner over the sequence of eUNG (Figure 6A – heat map). In simple words, it shows the per-residue *in vitro* evolution as a function of the specific bait. This analysis shows that the three populations evolved against purified GroEL, DnaKJ and CE-U cluster together, and are very different from the CE+U population in terms of positions positively enriched for mutations.

**Figure 6.**
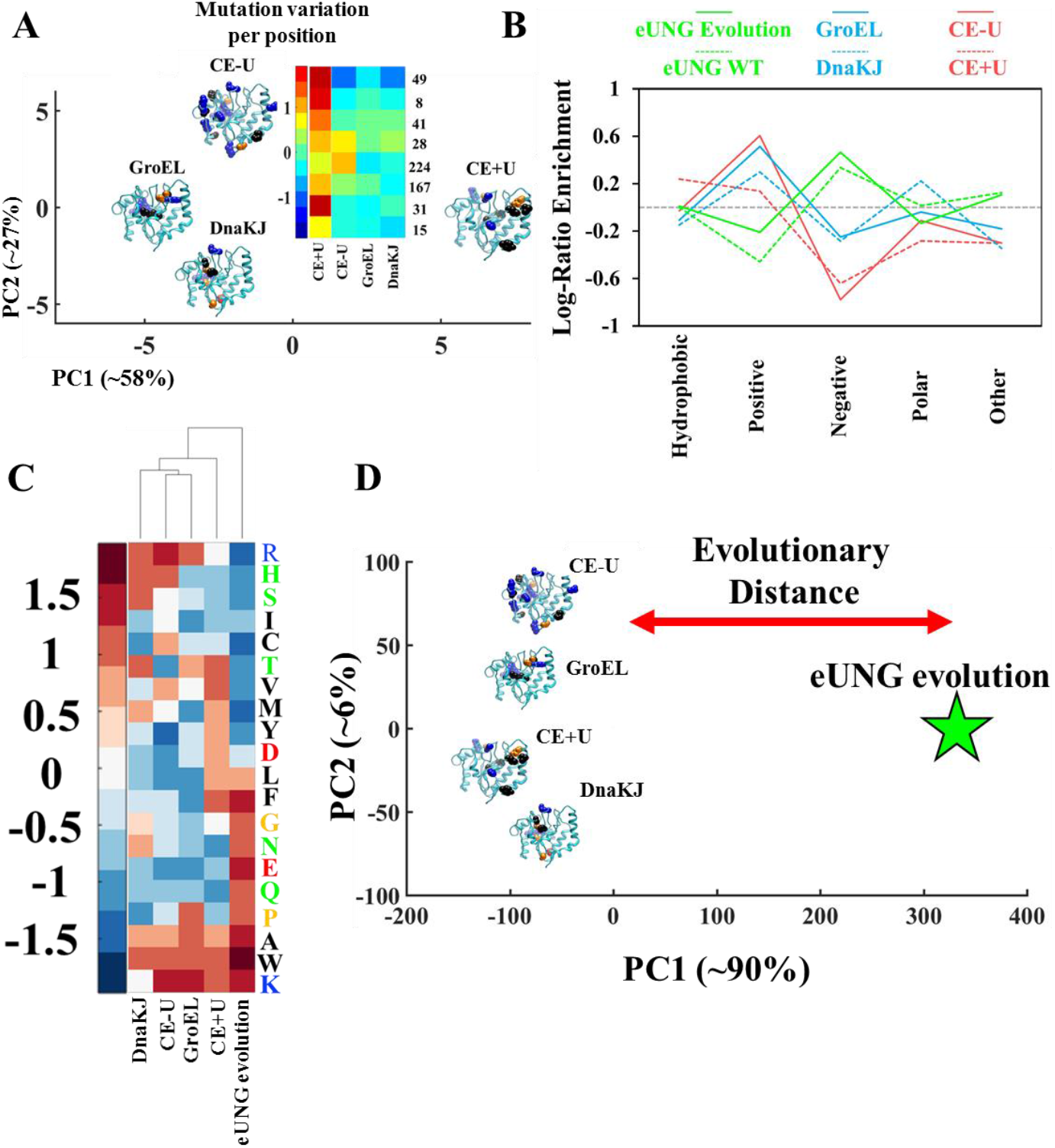
Covariance of enriched chaperones or CE sorted populations. **A.** PCA of the log-ratio values of DnaKJ, GroEL and CE+/-U binding populations. For each population positive log-ratio values were mapped on eUNG (PDB = 1FLZ) structure with their corresponding mutations. Axis percentage reflect the explained variance by a given principle component. PCA was calculated using the log ratio data derived using Eq. 1 (see materials and methods). Insert is the log-ratio clustering of enriched per position mutations in an area of interest of the cluster. Numbers on the right indicate variance position. Data is shown as a heat map. Each population log-ratio was combined to form an n-by-m matrix where n is the row number of the log-ratio in each position of the eUNG sequence and m is the chaperones/CE enriched populations. Red indicates high variance and blue low variance, per position. Values were standardized to have mean of 0 and SD of 1. **B.** Frequency of type of amino acid normalized by the theoretical frequency as calculated from the codon usage (Figure 3B and Table S4). The probability of having hydrophobic, positive, negative, polar or other side chain was calculated for selected populations. eUNG Evolution + WT are colored in green, chaperones with blue and cell extract binding populations with red. **C.** Sequence space of enriched GroEL, DnaKJ and CE+U binding populations as well as eUNG natural evolution (from ConSurf). The average frequency of each mutation, along the protein sequence, was normalized by the unsorted library (see Equation 3). The common log of these values was plotted. **D.** Co-variance of enriched population’s sequence space. Co-variance was calculated by PCA for the average frequency enrichment relative to the unsorted library (Equation 2 and 3), of each mutation that resulted in CE, GroEL and DnaKJ selection. Data was compared to ConSurf frequency matrix (Table S3). X axis resembles evolutionary distance (marked by two-sided arrow) where clustered dots suggest similar SP. Pentagram - ConSurf as evolutionary conservation. Analysis was done using Matlab R2017A).

Specific spatial distribution (SSD) was the second element that was thought to influence promiscuous evolution. It consists of amino acid residues, which were either negatively or positively selected relative to the starting library, which we have shown to represent random amino-acid frequencies in accordance to random single nucleotide mutations (therefore, the values in Figure 2B are around zero). Yet, as 8 codons code for positively charged residues and only 4 for negatively charged residues, the random library contains 13% Arg+Lys and only 6.5% Asp+Glu (Table S4), and thus the random protein is on average positively charged. Counting the amino-acid types in WT eUNG and averaging them over homologous UNG proteins as retrieved by ConSurf, shows frequencies that are richer in negatively charged residues and poorer in positively charge residues than the random library (Figure 6B), giving the protein a net charge of −2. This suggests evolutionary pressure towards a more neutral charge distribution of UNG than randomly expected, when starting from the eUNG sequence (Cohen-Khait and Schreiber, 2016). Next, we calculated the amino-acid frequencies of the different selected populations and grouped them according to positive, negative, polar, hydrophobic and other amino-acids (Figure 6B). Interestingly, GroEL, DnaKJ and CE-U selections resulted in further enrichment of positively charged amino-acids and fewer negatively charged amino-acids relative to eUNG-RL (which already has excess positive charge amino-acids), which is in contrast to natural UNG evolution (Figure 6B). CE+U resulted in fewer negatively charged residues together with fewer polar residues (Table S4), suggesting that evolutionary pressure towards cell extract binding while maintaining Ugi binding, is very different from that dictated by the fitness of a protein for its biological function (which includes avoiding the binding to other proteins in the cell extract). To obtain a more detailed picture, the log enrichment of the individual amino-acid mutations of each population was clustered and compared (Figure 6C). Strikingly, while both GroEL and DnaKJ-selected populations displayed an increase in positive charge, their amino acid distributions were different. Whereas both selected populations were enriched with arginines, only the GroEL population was enriched with lysines, which are more hydrophobic. In addition, GroEL selection favored prolines, while DnaKJ did not. Clustering shows CE-U to cluster closest to GroEL, in line with both being the most sensitive to protease cleavage. DnaKJ also cluster with the two other misfolded populations, while CE+U and eUNG evolution evolve differently from the misfolded populations and differently from one another. In quantitative terms, the variance between eUNG evolution and all selected populations is very large (PC1, Figure 6D), showing the large evolutionary pressure asserted by fitness (maintaining function, not just folding).

Using a more structural view on the locations of the most abundant mutations in the SSD, we mapped them on the eUNG structure. The log ratio values of the most abundant mutations of the chaperone-binding populations are mapped near the C-terminus (Figure S2), which, as a consequence, is more positively charged. To better understand the observed increased selection for positively-charged residues, observed for selection against chaperone proteins, we calculated the electrostatic potentials of eUNG and Ugi (Figure S3). Ugi shows a strong negatively-charged surface pointing towards eUNG, which displays a positively charged surface, albeit weaker, towards Ugi. Therefore, while the increased positive charge in chaperone-selected eUNG populations should benefit the protein-inhibitor complex binding, its lack suggests that it is more prone for misfolding.

## Discussion

### On the role of unfolding-chaperones in protein evolution

In vitro evolution of eUNG directed for chaperone or CE binding is very different from its natural variability, as depicted by ConSurf (Figure 6D). Natural evolution has a functional purpose, with selection directed towards keeping fitness and protein homeostasis, for which avoiding promiscuity is a major component. Evolving eUNG-RL against bacterial cell extract, resulted in yet two additional populations, with CE-U being close to GroEL and DnaKJ selected populations, while CE+U occupies another sequence space favoring cell extract binding. This population is limited only by foldability not by its fitness. We have observed that the CE-U selected population is closely related to the DnaKJ, and even more to the GroEL selected populations, but it also harbors unique properties. This can be explained by the existence of additional chaperones in CE, such as Tig, HtpG and ClpB (although note that CE selection was done without ATP). However, in the cell, selection is not necessarily driven towards chaperone-binding of misfolded states of proteins, and that chaperone binding can lead to degradation (Fernández-Fernández et al., 2017; Rizzolo et al., 2021). In this sense, our study reveals the different misfolded populations binding GroEL, DnaKJ or other chaperons (included in the CE-U population) and their fate upon trying to refold them as is discussed below.

Thermodynamically, our results imply that *in vitro* evolution towards chaperone-binding is accompanied by, and thus possibly because of, a loss of free energy and destabilization. However, as binding to chaperones leads eventually towards sorting the protein for degradation, as is the case of Hsp70 and proteasomal degradation (Fernández-Fernández et al., 2017), during the process of evolution, ATP-fueled unfoldase chaperones, by virtue of their ability to bind new, excessively unstable mutant proteins, may either “skim out” the excessively destabilizing mutations, or, as shown here with the complete active DnaK+DnaJ+GrpE+ATP system, give some unstable chaperone-binding mutant proteins another chance to natively refold (Figure 5C, for DnaKJ selected population, addition of KJE+ATP shows Ugi binding). This leads proteins to retain only near-native mutations that can either avoid chaperones or when they are misfolding, to be unfolded and refolded by them, until new stabilizing mutations are selected to evolve new functions (see Fig 7 above the dashed line).

**Figure 7.**
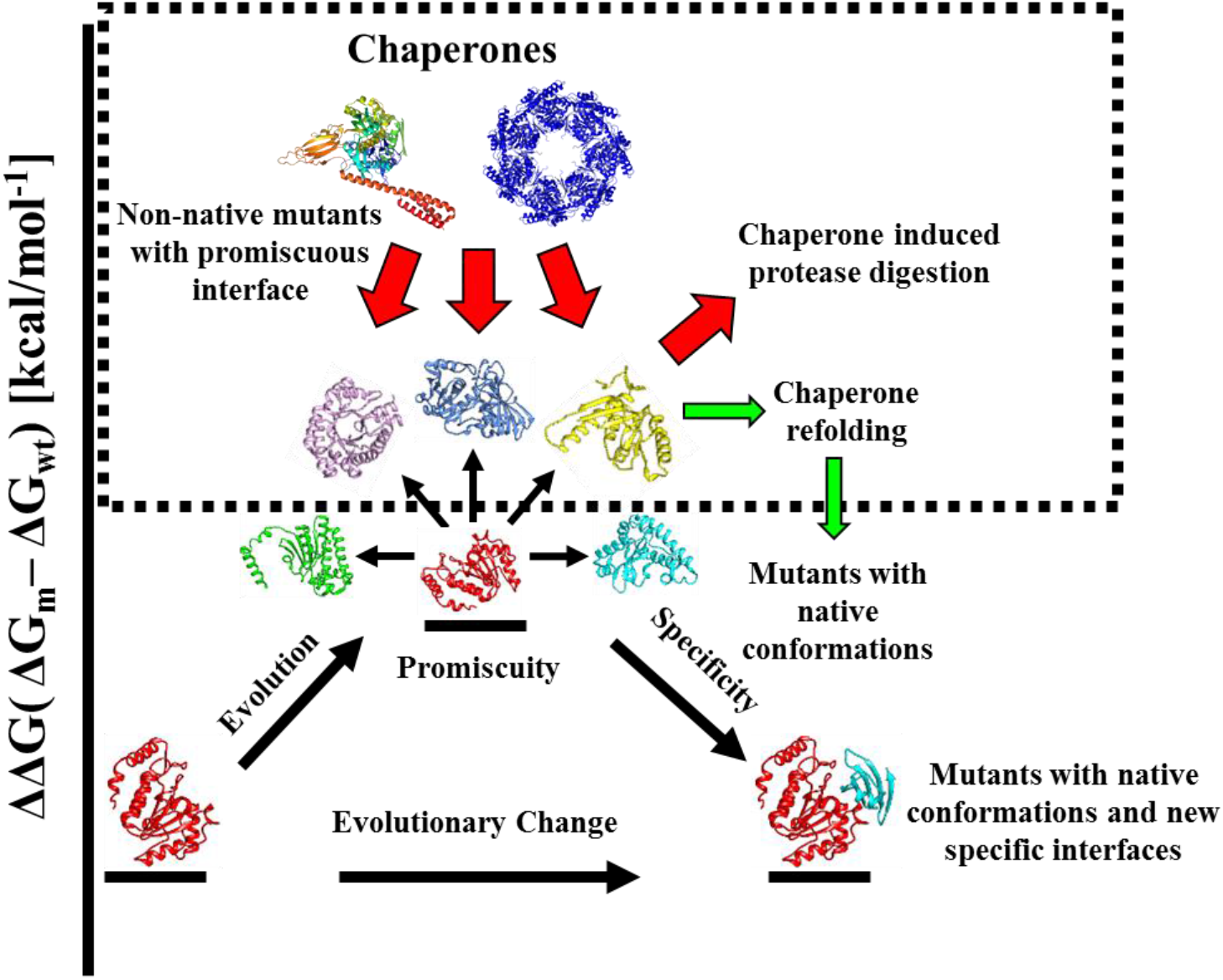
Chaperones modulate the evolutionary landscape of protein-protein interactions. Mutations in proteins can result in the formation of promiscuous interactions with non-target proteins. Chaperones promote evolution by buffering mutations, expanding the sequence space (SP) in which a protein can evolve. However, expanding the number of mutations a protein endures increases the likelihood it will mis-interact via misfolded species. While chaperones can maintain a folded state of a marginally destabilized conformations (green arrow), chaperones mostly selects against unstable promiscuous SPs (red arrows), promoting their degradation, until a near native solution is achieved, which is no longer recognized by chaperones.

Clear differences were observed between selection against pure GroEL_14_ oligomers (without ATP) and pure DnaK+DnaJ in the presence of ATP. These significant disparities motivated our reexamination of the different responses to partial protease digestion of these populations (Figure 5B). Since trypsin cleaves after lysine or arginine, and prolines leads to backbone breaks and local unfolding, their enrichment in the GroEL-selected sequence space suggests more disordered eUNG conformations, and thus higher protease sensitivity, compared to DnaKJ selected populations. Furthermore, the increase in positive charge and polarity in the GroEL- and DnaKJ-selected populations, resembles properties observed in intrinsically disordered proteins (IDPs) (Uversky, 2002). Particularly, the ratio between lysine and arginine was shown to influence the conformational equilibria of IDPs. While an increase in arginine over lysine in the DnaKJ-selected population promotes compactness in IDPs, higher lysine to arginine ratio in the GroEL-selected population promotes weaker phase-separation and results in reduced compactness of the misfolded species (Fossat et al., 2021; Sørensena and Kjaergaarda, 2019). This implies that the equilibria between unfolded, misfolded and native eUNG conformations, greatly differed in the GroEL, as compared to the DnaKJ-selected populations, enhancing susceptibility to protease digestion for the first. Increase in arginine residues might also lead to solubilization of the misfolded state, as arginine has been shown to enhance the solubility of aggregates or unstructured proteins (Austerberry et al., 2019). In addition, positive charges surrounding a hydrophobic core have been shown to characterize DnaK-binding motifs (Clerico et al., 2015; Rüdiger et al., 1997). As eUNG selection against DnaKJ resulted primarily in an increase in positive charges, it is plausible that this contributes to the observed increase in DnaK binding.

The protease degradation of selected populations (Figure 5B) might mimic *in vivo* evolution, because it may resemble what could have been the natural repartition of the work between ATP-fueled unfolding chaperones and proteases gated by ATP-fueled unfolding chaperones in evolution (Fauvet et al., 2021). In our assay, we differ from the conditions in the cell, as the proteins, chaperones and proteases are about tenfold less crowded than in cells. However, it is plausible that our *in vitro* chaperone selection is not so different than in the cell where the chaperones work hand in hand with proteases, and the non-promiscuous new mutants that are slightly less stable, may indeed be actively stabilized into native species, as previously been shown (Jarosz et al., 2010; Tokuriki and Tawfik, 2009). Those that are excessively less stable than the native WT, which cannot be stabilized to the native state by ATP-fueled unfolding/refolding chaperones may instead be hijacked by the AAA+ gated unfoldases/proteases that are in the same cellular compartment, and will be degraded and eliminated from evolution.

GroEL and DnaKJ chaperones recognize exposed motives at the surface of misfolded proteins that are mostly hydrophobic, which in the case of DnaK are also flanked by positive charges. Noticeably, these can be naturally enriched in mutants through a natural bias in evolution: the codon usage bias dictates that random mutagenesis should more frequently generate spontaneously sticky mutations, mainly hydrophobic and also positive charges that can interact with the negative charges that are often on the surface of native soluble proteins (Di Savino et al., 2021) (Figure 6A and B and Table S4). A protein can explore a larger mutational space and even include promiscuity-enhancing mutations due to the chaperones acting as enzymes that can iteratively unfold misfolded but not native protein conformations (Priya et al., 2013a). Whereas the codon usage bias increases the likelihood of promiscuity-enhancing mutations, chaperones have evolved to precisely detect this bias and apply a selection pressure against it (Buchberger et al., 1996; Rüdiger et al., 1997). Our results show that the chaperones will bind only to those mutants, as they indeed select against destabilizing mutations. This allows for the subsequent new stabilizing mutations, rendering the protein more native-like, i.e. that is not further recognized by chaperones (Rutherford, 2003; Tokuriki and Tawfik, 2009). The result is a feedback loop in which the component that permits in evolution the enhancement of the mutation load, also inhibits the possible enriched promiscuous products of this enhancement, by its destabilization. Inhibiting promiscuity suits well the chaperones, which can be 5-10% of a cell’s protein mass (Finka and Goloubinoff, 2013) and are therefore very expensive, but have value as major organizers of the cellular protein-protein interaction network (Calloni et al., 2012). If chaperones were lacking this feature, then protein stickiness should have been the null hypothesis (Aakre et al., 2015; Aharoni et al., 2005; Cohen-Khait et al., 2017; Hochberg et al., 2020; Schreiber and Keating, 2011). Any time scale for a solution to emerge would have to be found in a chaperone-dependent manner. Thus, along the tree of life, an ever increasing chaperone-addiction was necessary for the formation of the most complex proteomes, such as metazoan, plant and fungi proteomes (Rebeaud et al., 2021). It also explains why an organism complexity is not reflected by the number of distinct proteins it produces, but rather by the number of new folds (Taipale et al., 2014), the extend of fold combinations and of domain repeats resulting in increasingly long beta-fold enriched proteins, as well as of chaperone-indifferent IDPs, of which some that form toxic amyloids are also chaperone-resistant, along evolution (Rebeaud et al., 2021). Noticeably, our results suggest that the IDPs that did survive chaperone selection in evolution would have to lack typical chaperone-binding motives, such as clusters of hydrophobic residues, flanked by positive charges, as identified before and here (Goloubinoff et al., 1989; Jakob et al., 1993; Koldewey et al., 2016; Rüdiger et al., 1997). Furthermore, it might explain why core-chaperones have not evolved proportionally with the increased complexity of proteomes. Rather co-chaperones, such as the J-domain proteins, of which DnaJ is a member, which specifically target the Hsp70 unfoldase machineries to bind misfolded protein substrates with high-affinity and use energy of ATP to unfold them. This, while avoiding binding to the natively unfolded and the compact native proteins, which are in general the low-affinity products of the chaperone-mediated unfolding reaction (Rebeaud et al., 2021). In addition, our results suggest that the Hsp60 and Hsp70 chaperones, alongside being ATP-fueled unfoldases, may also assist the proper assembly of protein oligomers. Chaperone assisted assembly was first suggested by John Ellis and colleagues, who showed that the chloroplast HSP60 (CPN60) was key to the proper assembly of RubisCO L_8_S_8_ dodecamers (Hemmingsen et al., 1988). It seems that the term chaperones, initially chosen by John Ellis to describe features of this class of proteins, was not inaccurate. As such, chaperones are required for specific protein multimerization, thus it is plausible that they inhibit promiscuous interactions, and allow the formation only of rare specific interactions, in particular native oligomers and thereby limiting their evolvability (Duraõ et al., 2015). In conclusion, we suggest that the evolutionary landscape of protein promiscuity is modulated by molecular chaperones, performing as the molecular “police” of cells (Hinault and Goloubinoff, 2007) to shield promiscuous, counter-productive protein-protein interactions, carried out primarily by misfolding and aggregating proteins, which are cytotoxic and cause degenerative diseases in metazoans. This passively drives specificity, as only binding interfaces, which are not recognized by chaperones are positively selected for, in the course of protein evolution (Figure 7).

## Materials and Methods

### Yeast library formation

A library of randomly mutated *E.coli* Uracil glycosylase (eUNG) was created on EBY100 yeast by following the procedure described by Benatuil (Benatuil et al., 2010). DNA mutagenesis was performed with mutazyme (GeneMorph II random mutagenesis kit catalog # 200550) resulting in an average of ~4 mutations per gene. Amplification of eUNG gene was done with Taq DNA-polymerase as described by Chao (Chao et al., 2006). Library size was estimated to be ~10^7^ by plating serial dilutions on YPD selection plates lacking Tryptophan. Library was further confirmed for correct gene insertion and average mutation by sequencing 20 single clones.

### Co-incubation experiments

After final round of selection against chaperones or WT cell extract, enriched populations were collected and incubated both with chaperones or WT cell extract and Ugi. Chaperones and WT cell extracts were labeled with APC (Biotium # 92108) while Ugi was labeled with FITC (Biotium # 92103) as described by the manufacturer. Co-incubated populations were analyzed by Bio-Rad FACS sorter and according to population distribution were sorted out into new vials with SDCCA selection media (Chao et al., 2006).

### Uracil glycosylase inhibitor production

Urcail glycosylase inhibitor (Ugi) gene was ordered from TWIST and later cloned into the pet28 vector.(Zahradník et al., 2019) The vector was modified to have a longer linker on the N’ terminus for better protease cleavage. Ugi was expressed in BL21(DE3) bacteria, grown at 37 ° C until O.D_600_ = 0.6-0.8 and then incubated with 0.1 mM IPTG at 16 ° C overnight. Bacteria were pelleted and sonicated and suspended in SUMO protease buffer (50mM Tris-HCL pH 8, 200mM NaCl 10% glycerol 1mM DTT and 1:200 sumo protease both of which were added fresh). Ugi was purified from Ni beads binding 6xHis tag as described in (Zahradník et al., 2019) and further purified using ion exchange chromatography. After cleavage Ugi was dialyzed into labeling buffer (0.1M NaHCO_3_ pH = ~8.3).

### Cell extracts production

W3110 *E.coli* WT cell extract was prepared by first inoculating at 1:1000 ratio into 2YT followed by incubation at 37 ° C until O.D_600_ of 0.6-0.8. Cells were then harvested by centrifugation at 4 ° C and resuspended in lysis buffer (Tris HCl pH = 8, 200 mM NaCl, 10 mM Imidazole and 10% glycerol). The bacteria were sonicated and cytosolic fraction was collected by 17000 rpm 4 ° C for 30 minutes. Supernatant was filtered using 0.4 μM filters and was dialyzed in labeling buffer (0.1M NaHCO_3_ pH = ~8.3) in double distilled water and in 3.5 MWCO dialysis membrane.

### Yeast library sorting

The library was sorted 5 times against 5 mg/ml cell extract labeled with CF640R succinimidyl ester (APC) through amine-coupling. APC labeled cell extract was filtered from excess dye before each sort. Protein expression levels on the yeast surface was detected by labeling the c-Myc tag found at the C’ terminal of expressed eUNG with FITC coloring. The selection continued until notable binding enrichment was detected by FACS. The selected libraries were then plated and 20 clones were sequenced and evaluated for binding cell extracts or chaperones.

### Deep Sequencing

Deep sequencing was done on all the populations. Plasmids encoding the selected populations were extracted using Lyticase (Sigma # L2524). The eUNG gene (636bp) was amplified as 4 fragments of 250 bp each. Amplified plasmid flanks, which are not part of the eUNG gene were discarded in the analysis. Deep sequencing was done using Illumina MiSeq V2 (500 cycles) with paired-ends, 250bp read length and 12M reads. Deep sequencing reads were extracted using Enrich2 (Rubin et al., 2017) and were further processed using Matlab R2017a. Sequence logos images were done using Seq2Logo (Thomsen and Nielsen, 2012).

### Matlab analysis of deep sequencing data

Deep sequencing raw data was extract using Enrich2. The reads were organized in excel files containing reads per mutation along eUNG sequence. Reads were used to write scripts (Extended data) which calculates three parameters; Log-Ratio – SD (Eq. 1), amino acid frequency per position and from it the SSD (Eq. 2 and 3) and the initial amino acid variance, denoted by V_AA_, of the unsorted library (Eq. 4):

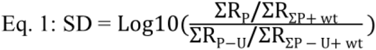

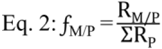

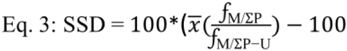

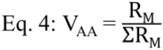

ΣR_P_ – total reads per position, ΣR_∑P + WT_ – total reads over all positions including WT sequences, ΣR_P-U_ – unsorted library total reads per position, ΣR_∑P-U + WT_ – unsorted library total reads over all positions including WT sequences, 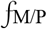 – mutation frequency per position, R_M/P_ – reads of a mutation per position, 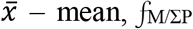 – frequency of a mutation over all positions. 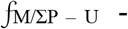 unsorted library frequency of a mutation over all positions, V_AA_ – initial amino acid variance (the frequency of a possible mutation for an amino acid) displayed as a fraction of: R_M_ – reads of a specific mutation X of amino acid Y and ΣR_M_ – sum of all reads of possible mutations for an amino acid Y.

## Acknowledgement

We thank Satyam Tiwari and Tatiana Fomekong T. Mbefo from the University of Lausanne (UNIL), for providing us with purified GroEL, DnaK and DnaJ proteins and Pierre Gevevaux from the University of Toulouse III for giving us the delta DnaJK-DnaJ *E. coli* stain.

## Supporting data

**Figure S1.**
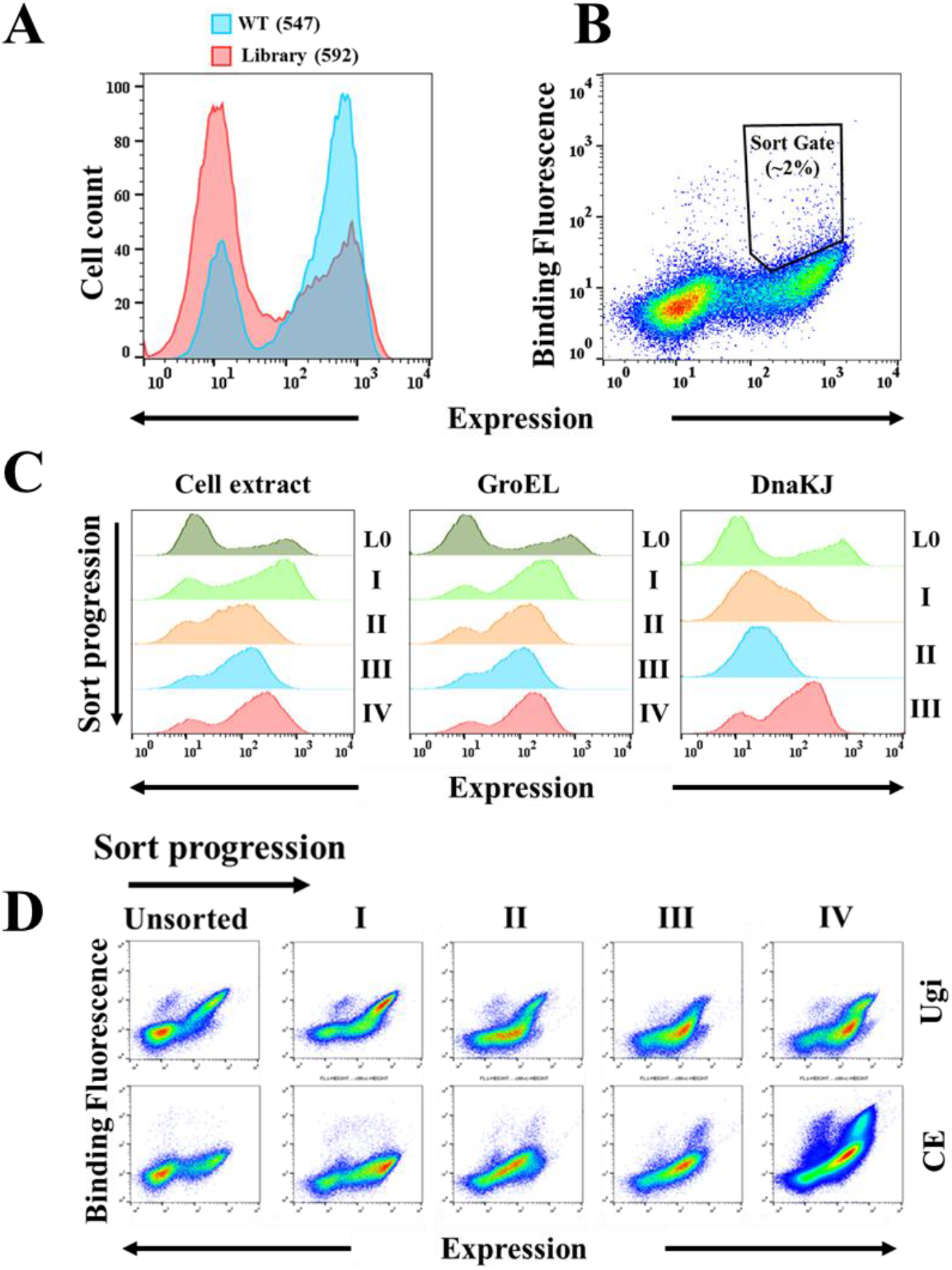
Binding and expression profile of eUNG library. **A.** Expression distribution of eUNG WT compared with eUNG random library. Both eUNG WT and library were labeled for expression using anti-cMyc primary AB (1:50 ratio) and after with FITC secondary AB (1:100 ratio). Legend numbers indicate the mean fluorescence value for WT (light blue) and library (red). **B.** Illustration of sort gate used in selection round to chaperones and cell extract. Selection was done such that only 2% of expressing population was sorted. In the first round of selection ~10^7^ samples were sorted to cover the entire genetic complexity of the library. **C.** Expression of eUNG library as a function of selection rounds and a specific bait (cell extract, GroEL and DnaKJ). L0 indicates the unsorted library. **D.** Loss of Ugi binding as a function of cell extract binding enrichment. Top and bottom panels show Ugi and cell extract binding at each round of selection, respectively. For both C&D, each round of selection is marked with roman numerals and sort progression with a black arrow. Data was analyzed using FlowJo.

**Figure S2.**
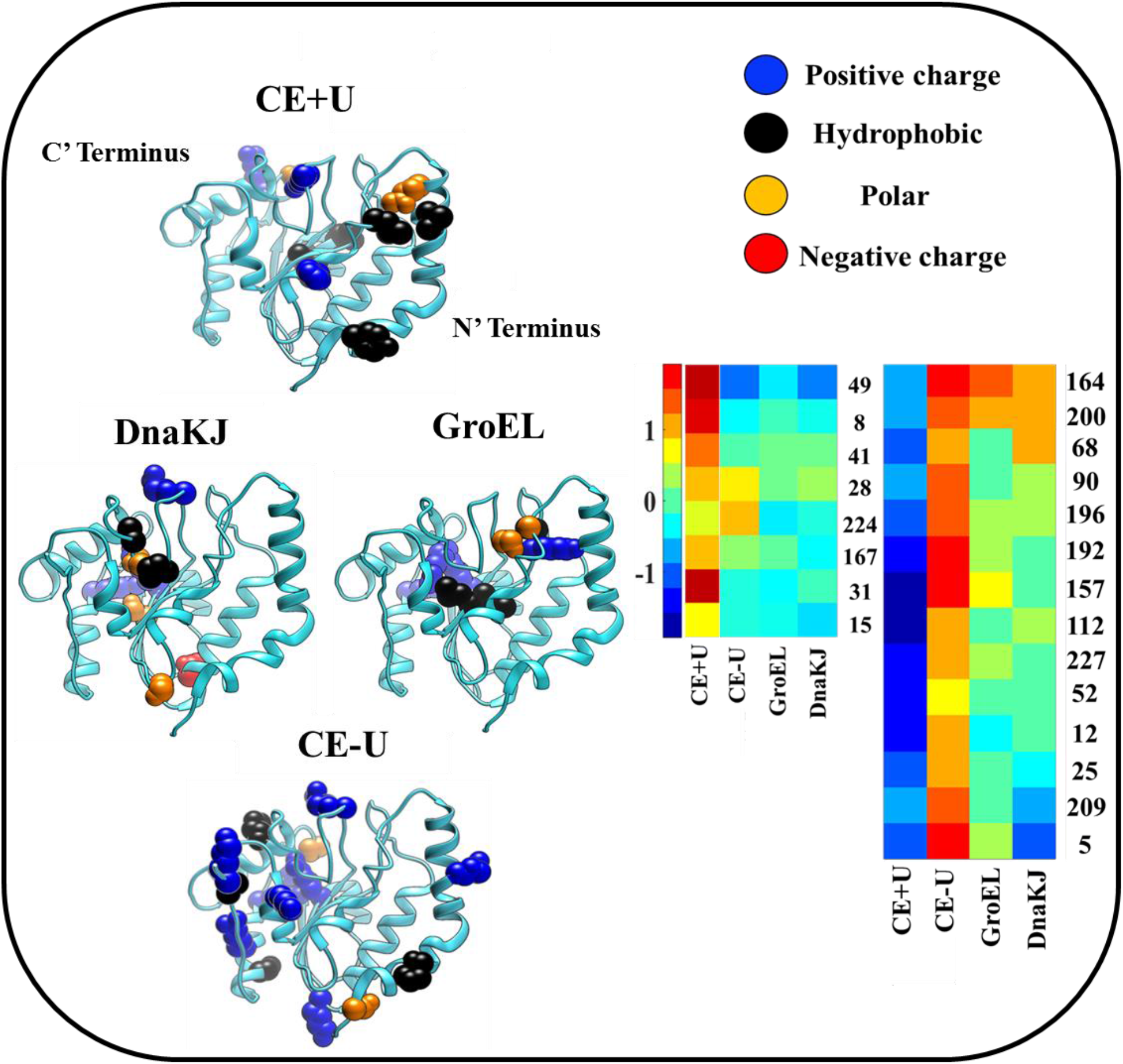
Structural mapping of positions positively selected for binding. Positions that were enriched five-fold compared to the unsorted library were mapped on UNG (PDB: 1EUG). Each position was mutated to its most abundant residue according to each log ratio value (Table S1) and colored according to its chemical property. Black – hydrophobic, blue – positive charge, orange – polar and red – negative charge. Insert is the log-ratio clustering of enriched per position mutations in an area of interest of the cluster. Numbers on the right indicate variance position. Data is shown as a heat map. Each population log-ratio was combined to form an n-by-m matrix where n is the row number of the log-ratio in each position of the eUNG sequence and m is the chaperones/CE enriched populations. Red indicates high variance and blue low variance, per position. Values were standardized to have mean of 0 and SD of 1. Structural analysis was done using Chimera.

**Figure S3.**
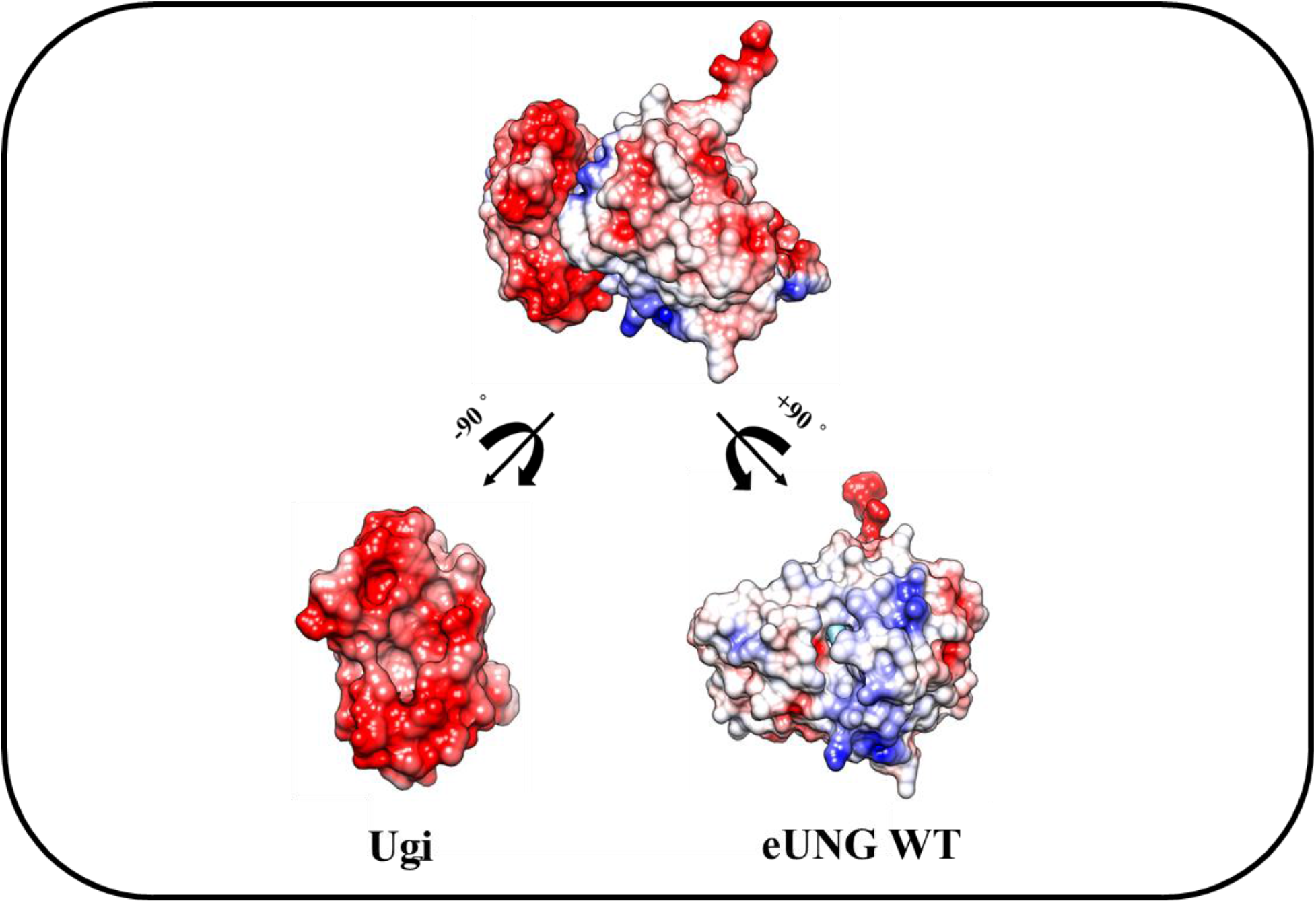
eUNG WT and Ugi interact via complementary charge interface. eUNG WT and Ugi in complex along with -/+90 ° shift for Ugi and eUNG WT, respectively. Red indicates negative charge and blue positive. White color are non-charged residues. Electrostatic potential was calculated using Chimera.

**Table S4.**
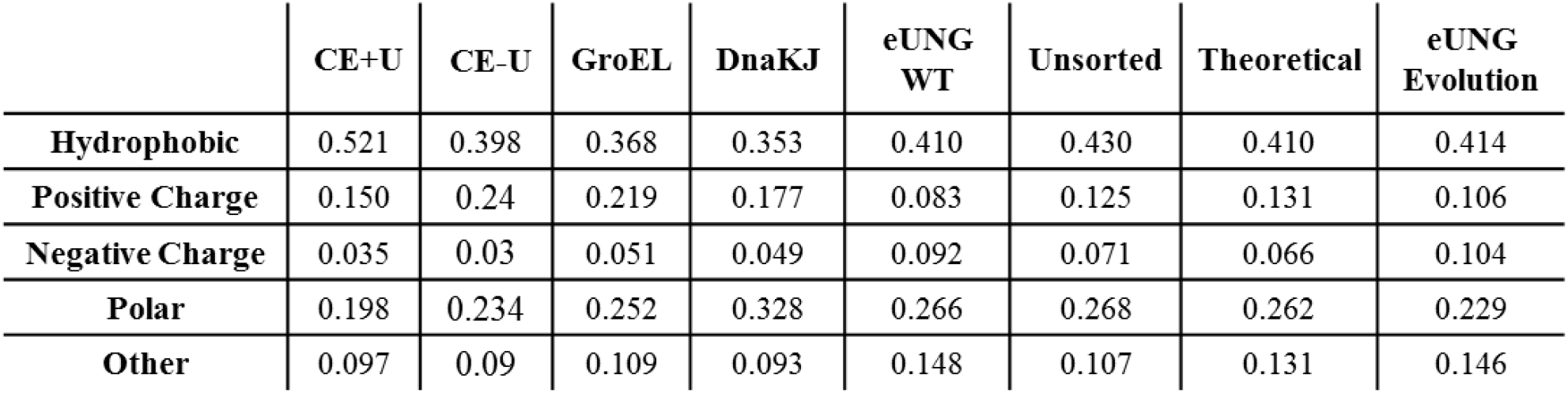
Amino acid type frequency matrix. The fraction of each amino acid type was calculated by taking the average of sums for each hydrophobic, positive charge, negative charge and other, amino acids. The theoretical fractions represents that number of codons for a certain amino acid type compared to the whole codon usage. Amino acid type includes: Also, eUNG evolution frequencies were extracted out of ConSurf mutation frequency matrix. Hydrophobic (LVIWYFACM), Positive charge (RK), Negative charge (DE), Polar (NQSTH) and other (PG). The natural log ratio between these propensities (compared to the theoretical) were used to calculate the log enrichments of Figure 3B and Figure 6B.

|  | **CE+U** | **CE-U** | **GroEL** | **DnaKJ** | **eUNG**<br>**WT** | **Unsorted** | **Theoretical** | **eUNG**<br>**Evolution** |
| --- | --- | --- | --- | --- | --- | --- | --- | --- |
| **Hydrophobic** | 0.521 | 0.398 | 0.368 | 0.353 | 0.410 | 0.430 | 0.410 | 0.414 |
| **Positive Charge** | 0.150 | 0.24 | 0.219 | 0.177 | 0.083 | 0.125 | 0.131 | 0.106 |
| **Negative Charge** | 0.035 | 0.03 | 0.051 | 0.049 | 0.092 | 0.071 | 0.066 | 0.104 |
| **Polar** | 0.198 | 0.234 | 0.252 | 0.328 | 0.266 | 0.268 | 0.262 | 0.229 |
| **Other** | 0.097 | 0.09 | 0.109 | 0.093 | 0.148 | 0.107 | 0.131 | 0.146 |

## References

1. Hartl FU, Bracher A, Hayer-Hartl M. 2011. Molecular chaperones in protein folding and proteostasis. Nature 475:324–332. doi:10.1038/nature10317

2. Goloubinoff P, Sassi AS, Fauvet B, Barducci A, De Los Rios P. 2018. Chaperones convert the energy from ATP into the nonequilibrium stabilization of native proteins article. Nat Chem Biol 14:388–395. doi:10.1038/s41589-018-0013-8

3. Finka A, Mattoo RUH, Goloubinoff P. 2016. Experimental Milestones in the Discovery of Molecular Chaperones as Polypeptide Unfolding Enzymes. Annu Rev Biochem 85:715–742. doi:10.1146/annurev-biochem-060815-014124

4. Rebeaud ME, Mallik S, Goloubinoff P, Tawfik DS. 2021. On the evolution of chaperones and cochaperones and the expansion of proteomes across the Tree of Life. Proc Natl Acad Sci U S A 118. doi:10.1073/pnas.2020885118

5. Tokuriki N, Tawfik DS. 2009. Chaperonin overexpression promotes genetic variation and enzyme evolution. Nature 459:668–673. doi:10.1038/nature08009

6. Libich DS, Tugarinov V, Clore GM. 2015. Intrinsic unfoldase/foldase activity of the chaperonin GroEL directly demonstrated using multinuclear relaxation-based NMR. Proc Natl Acad Sci U S A 112:8817–8823. doi:10.1073/pnas.1510083112

7. Agozzino L, Dill KA. 2018. Protein evolution speed depends on its stability and abundance and on chaperone concentrations. Proc Natl Acad Sci U S A 115:9092–9097. doi:10.1073/pnas.1810194115

8. Rutherford SL. 2003. Between genotype and phenotype: Protein chaperones and evolvability. Nat Rev Genet 4:263–274. doi:10.1038/nrg1041

9. Rüdiger S, Schneider-Mergener J, Bukau B. 2001. Its substrate specificity characterizes the DnaJ co-chaperone as a scanning factor for the DnaK chaperone. EMBO J 20:1042–1050. doi:10.1093/emboj/20.5.1042

10. Rüdiger S, Buchberger A, Bukau B. 1997. Interaction of Hsp70 chaperones with substrates. Nat Struct Biol 4:342–349. doi:10.1038/nsb0597-342

11. Horovitz A, Reingewertz TH, Cuéllar J, Valpuesta JM. 2022. Chaperonin Mechanisms: Multiple and (Mis)Understood? 115–133.

12. Priya S, Sharma SK, Sood V, Mattoo RUH, Finka A, Azem A, De Los Rios P, Goloubinoff P. 2013. GroEL and CCT are catalytic unfoldases mediating out-of-cage polypeptide refolding without ATP. Proc Natl Acad Sci U S A 110:7199–7204. doi:10.1073/pnas.1219867110

13. Stiffler MA, Hekstra DR, Ranganathan R. 2015. Evolvability as a Function of Purifying Selection in TEM-1 β-Lactamase. Cell 160:882–892. doi:10.1016/j.cell.2015.01.035

14. Hochberg GKA, Liu Y, Marklund EG, Metzger BPH, Laganowsky A, Thornton JW. 2020. A hydrophobic ratchet entrenches molecular complexes. Nature 588:503–508. doi:10.1038/s41586-020-3021-2

15. De Los Rios P, Barducci A. 2014. Hsp70 chaperones are non-equilibrium machines that achieve ultra-affinity by energy consumption. Elife 3:1–10. doi:10.7554/elife.02218

16. Sekhar A, Velyvis A, Zoltsman G, Rosenzweig R, Bouvignies G, Kay LE. 2018. Conserved conformational selection mechanism of Hsp70 chaperone-substrate interactions. Elife 7:1–29. doi:10.7554/eLife.32764

17. Rosenzweig R, Sekhar A, Nagesh J, Kay LE. 2017. Promiscuous binding by Hsp70 results in conformational heterogeneity and fuzzy chaperone-substrate ensembles. Elife 6:1–22. doi:10.7554/eLife.28030

18. Pearl LH. 2000. Structure and function in the uracil-DNA glycosylase superfamily. Mutat Res - DNA Repair 460:165–181. doi:10.1016/S0921-8777(00)00025-2

19. Rutherford SL, Lindquist S. 1998. Hsp90 as a capacitor for morphological evolution. Nature 396:336–342. doi:10.1038/24550

20. Buchberger A, Schröder H, Hesterkamp T, Schönfeld HJ, Bukau B. 1996. Substrate shuttling between the DnaK and GroEL systems indicates a chaperone network promoting protein folding. J Mol Biol 261:328–333. doi:10.1006/jmbi.1996.0465

21. Worth CL, Preissner R, Blundell TL. 2011. SDM - A server for predicting effects of mutations on protein stability and malfunction. Nucleic Acids Res 39:215–222. doi:10.1093/nar/gkr363

22. Tokuriki N, Stricher F, Schymkowitz J, Serrano L, Tawfik DS. 2007. The Stability Effects of Protein Mutations Appear to be Universally Distributed. J Mol Biol 369:1318–1332. doi:10.1016/j.jmb.2007.03.069

23. Biol M, Marel V Der, Boom V, Marel V Der, Biol CAJM, Lim WA, Farruggio DC, Sauer RT. 1992. Disruptive Mutations in * 4324–4333.

24. Krissinel E, Henrick K. 2007. Inference of Macromolecular Assemblies from Crystalline State. J Mol Biol 372:774–797. doi:10.1016/j.jmb.2007.05.022

25. Ashkenazy H, Abadi S, Martz E, Chay O, Mayrose I, Pupko T, Ben-Tal N. 2016. ConSurf 2016: an improved methodology to estimate and visualize evolutionary conservation in macromolecules. Nucleic Acids Res 44:W344–W350. doi:10.1093/nar/gkw408

26. Walsh KA, Kauffman DL, Kumar KS, Neurath H. 1964. on the Structure and Function of Bovine Trypsinogen and Trypsin. Proc Natl Acad Sci United States 51:301–308. doi:10.1073/pnas.51.2.301

27. Ebeling W, Hennrich N, Klockow M, Metz H, Orth HD, Lang H. 1974. Proteinase K from Tritirachium album Limber. Eur J Biochem 47:91–97. doi:10.1111/j.1432-1033.1974.tb03671.x

28. Priya S, Sharma SK, Goloubinoff P. 2013. Molecular chaperones as enzymes that catalytically unfold misfolded polypeptides. FEBS Lett 587:1981–1987. doi:10.1016/j.febslet.2013.05.014

29. Sharma SK, De Los Rios P, Christen P, Lustig A, Goloubinoff P. 2010. The kinetic parameters and energy cost of the Hsp70 chaperone as a polypeptide unfoldase. Nat Chem Biol 6:914–920. doi:10.1038/nchembio.455

30. Ben-Zvi A, De Los Rios P, Dietler G, Goloubinoff P. 2004. Active solubilization and refolding of stable protein aggregates by cooperative unfolding action of individual Hsp70 chaperones. J Biol Chem 279:37298–37303. doi:10.1074/jbc.M405627200

31. Fauvet B, Rebeaud ME, Tiwari S, De Los Rios P, Goloubinoff P. 2021. Repair or Degrade: the Thermodynamic Dilemma of Cellular Protein Quality-Control. Front Mol Biosci 8:1–11. doi:10.3389/fmolb.2021.768888

32. Sharma SK, De Los Rios P, Goloubinoff P. 2011. Probing the different chaperone activities of the bacterial HSP70-HSP40 system using a thermolabile luciferase substrate. Proteins Struct Funct Bioinforma 79:1991–1998. doi:10.1002/prot.23024

33. Goloubinoff P, Christeller JT, Gatenby AA, Lorimer GH. 1989. Reconstitution of active dimeric ribulose bisphosphate carboxylase from an unfolded state depends on two chaperonin proteins and Mg-ATP. Nature 342:884–889. doi:10.1038/342884a0

34. Diamant S, Peres Ben-Zvi A, Bukau B, Goloubinoff P. 2000. Size-dependent disaggregation of stable protein aggregates by the DnaK chaperone machinery. J Biol Chem 275:21107–21113. doi:10.1074/jbc.M001293200

35. Cohen-Khait R, Schreiber G. 2016. Low-stringency selection of TEM1 for BLIP shows interface plasticity and selection for faster binders. Proc Natl Acad Sci U S A 113:14982–14987. doi:10.1073/pnas.1613122113

36. Fernández-Fernández MR, Gragera M, Ochoa-Ibarrola L, Quintana-Gallardo L, Valpuesta JM. 2017. Hsp70 – a master regulator in protein degradation. FEBS Lett 591:2648–2660. doi:10.1002/1873-3468.12751

37. Rizzolo K, Yu AYH, Ologbenla A, Kim SR, Zhu H, Ishimori K, Thibault G, Leung E, Zhang YW, Teng M, Haniszewski M, Miah N, Phanse S, Minic Z, Lee S, Caballero JD, Babu M, Tsai FTF, Saio T, Houry WA. 2021. Functional cooperativity between the trigger factor chaperone and the ClpXP proteolytic complex. Nat Commun 12:1–18. doi:10.1038/s41467-020-20553-x

38. Uversky VN. 2002. What does it mean to be natively unfolded? Eur J Biochem 269:2–12. doi:10.1046/j.0014-2956.2001.02649.x

39. Sørensena CS, Kjaergaarda M. 2019. Effective concentrations enforced by intrinsically disordered linkers are governed by polymer physics. Proc Natl Acad Sci U S A 116:23124–23131. doi:10.1073/pnas.1904813116

40. Fossat MJ, Zeng X, Pappu R V. 2021. Uncovering Differences in Hydration Free Energies and Structures for Model Compound Mimics of Charged Side Chains of Amino Acids. J Phys Chem B 125:4148–4161. doi:10.1021/acs.jpcb.1c01073

41. Austerberry JI, Thistlethwaite A, Fisher K, Golovanov AP, Pluen A, Esfandiary R, Van Der Walle CF, Warwicker J, Derrick JP, Curtis R. 2019. Arginine to Lysine Mutations Increase the Aggregation Stability of a Single-Chain Variable Fragment through Unfolded-State Interactions. Biochemistry 58:3413–3421. doi:10.1021/acs.biochem.9b00367

42. Clerico EM, Tilitsky JM, Meng W, Gierasch LM. 2015. How Hsp70 molecular machines interact with their substrates to mediate diverse physiological functions. J Mol Biol 427:1575–1588. doi:10.1016/j.jmb.2015.02.004

43. Jarosz DF, Taipale M, Lindquist S. 2010. Protein homeostasis and the phenotypic manifestation of genetic diversity: Principles and mechanisms. Annu Rev Genet 44:189–216. doi:10.1146/annurev.genet.40.110405.090412

44. Di Savino A, Foerster JM, Ullmann GM, Ubbink M. 2021. The Charge Distribution on a Protein Surface Determines Whether Productive or Futile Encounter Complexes Are Formed. Biochemistry 60:747–755. doi:10.1021/acs.biochem.1c00021

45. Finka A, Goloubinoff P. 2013. Proteomic data from human cell cultures refine mechanisms of chaperone-mediated protein homeostasis. Cell Stress Chaperones 18:591–605. doi:10.1007/s12192-013-0413-3

46. Calloni G, Chen T, Schermann SM, Chang HC, Genevaux P, Agostini F, Tartaglia GG, Hayer-Hartl M, Hartl FU. 2012. DnaK Functions as a Central Hub in the E. coli Chaperone Network. Cell Rep 1:251–264. doi:10.1016/j.celrep.2011.12.007

47. Schreiber G, Keating AE. 2011. Protein binding specificity versus promiscuity. Curr Opin Struct Biol 21:50–61. doi:10.1016/j.sbi.2010.10.002

48. Aharoni A, Gaidukov L, Khersonsky O, Gould SMQ, Roodveldt C, Tawfik DS. 2005. The “evolvability” of promiscuous protein functions. Nat Genet 37:73–76. doi:10.1038/ng1482

49. Aakre CD, Herrou J, Phung TN, Perchuk BS, Crosson S, Laub MT. 2015. Evolving New Protein-Protein Interaction Specificity through Promiscuous Intermediates. Cell 163:594–606. doi:10.1016/j.cell.2015.09.055

50. Cohen-Khait R, Dym O, Hamer-Rogotner S, Schreiber G. 2017. Promiscuous Protein Binding as a Function of Protein Stability. Structure 25:1867–1874.e3. doi:10.1016/j.str.2017.11.002

51. Taipale M, Tucker G, Peng J, Krykbaeva I, Lin ZY, Larsen B, Choi H, Berger B, Gingras AC, Lindquist S. 2014. A quantitative chaperone interaction network reveals the architecture of cellular protein homeostasis pathways. Cell 158:434–448. doi:10.1016/j.cell.2014.05.039

52. Koldewey P, Stull F, Horowitz S, Martin R, Bardwell JCA. 2016. Forces Driving Chaperone Action. Cell 166:369–379. doi:10.1016/j.cell.2016.05.054

53. Jakob U, Gaestel M, Engel K, Buchner J. 1993. Small heat shock proteins are molecular chaperones. J Biol Chem 268:1517–1520. doi:10.1016/s0021-9258(18)53882-5

54. Hemmingsen SM, Woolford C, Van Der Vies SM, Tilly K, Dennis DT, Georgopoulos CP, Hendrix RW, Ellis RJ. 1988. Homologous plant and bacterial proteins chaperone oligomeric protein assembly. Nature 333:330–334. doi:10.1038/333330a0

55. Duraõ P, Aigner H, Nagy P, Mueller-Cajar O, Hartl FU, Hayer-Hartl M. 2015. Opposing effects of folding and assembly chaperones on evolvability of Rubisco. Nat Chem Biol 11:148–155. doi:10.1038/nchembio.1715

56. Hinault M-P, Goloubinoff P. 2007. Molecular Crime and Cellular Punishment. Mol Asp Stress Response Chaperones, Membr Networks 47–54. doi:10.1007/978-0-387-39975-1_5

